# ST6Gal2 promotes α2,6-sialylation and aggressive phenotypes in neuroblastoma cells

**DOI:** 10.64898/2026.09.24.754123

**Authors:** Laxmi Swami, Punarvi Mandadapu, Mayra E. Lopez, Sarah K. Vaughan, Abigail R. Richardson, Pracheta Janmeda, Karin M. Hardiman, Laura L. Stafman

## Abstract

Neuroblastoma is the most common extracranial solid tumor of childhood. Its clinical behavior ranges from spontaneous regression to lethal, treatment-refractory disease. Aberrant α2,6-sialylation contributes to aggressive phenotypes in many cancers, but the role of ST6Gal2, a neural-enriched α2,6-sialyltransferase, in neuroblastoma is largely unexplored. Here, we examine the clinical and functional significance of ST6Gal2 in neuroblastoma. In two independent public cohorts (SEQC, n=498; Kocak, n=649), high *ST6GAL2* expression was associated with significantly worse overall and event-free survival. In the SEQC cohort, *ST6GAL2* expression was higher in high-risk and *MYCN*-amplified tumors, varied across International Neuroblastoma Staging System stages, and correlated positively with a mesenchymal transcriptional signature (Spearman ρ=0.181). The mesenchymal correlation was reproduced in the Kocak cohort (ρ=0.204). Stable shRNA-mediated knockdown of *ST6GAL2* in SK-N-AS and SK-N-BE(2) cells reduced proliferation and viability, impaired wound closure, and decreased migration and invasion. In preliminary experiments in SK-N-AS cells, *ST6GAL2* knockdown reduced binding of *Sambucus nigra* agglutinin, consistent with a role for ST6Gal2 in α2,6-sialylation. Together, these findings link ST6Gal2 expression to aggressive clinical and transcriptional features and pro-tumorigenic phenotypes in neuroblastoma and nominate ST6Gal2-mediated sialylation as a candidate pathway for mechanistic study.

## Introduction

Neuroblastoma is the most common extracranial solid tumor of childhood and accounts for a disproportionate number of pediatric cancer deaths.^1^ Clinical behavior is highly heterogeneous and ranges from disease that undergoes spontaneous regression or differentiation to aggressive disease that persists despite intensive multimodal therapy.^2, 3^ Outcomes remain poor for patients with high-risk or relapsed neuroblastoma,^4^ highlighting the need to better define molecular pathways associated with aggressive tumor behavior so that we may better understand this disease and potentially develop novel therapeutics. Neuroblastoma cells exhibit distinct adrenergic and mesenchymal transcriptional states, with the mesenchymal state associated with aggressive features including increased invasiveness and therapeutic resistance.^5^

Aberrant glycosylation is increasingly recognized as an important feature of cancer biology.^6^ Sialylation, a particular form of glycosylation defined by the addition of sialic acid residues to terminal positions on glycoproteins, can alter receptor signaling, ligand binding, protein stability, and cell-cell interactions and thereby influence tumor cell behavior.^7^ α2,6-sialylation is catalyzed by the sialyltransferases ST6Gal1 and ST6Gal2.^8^ ST6Gal1 is broadly expressed and has been implicated in malignant phenotypes including cell survival, invasion, plasticity, and therapeutic resistance in adult cancers.^9-11^ In contrast, considerably less is known about the role of ST6Gal2 in cancer.

ST6Gal2 differs from ST6Gal1 in tissue distribution as it is enriched in neural tissues.^8^ Previous studies characterized *ST6GAL2* transcription and ST6Gal2 protein expression in neural tissues and in the SH-SY5Y neuroblastoma cell line, suggesting relevance in neural-derived malignancies such as neuroblastoma.^12^ Recently, *ST6GAL2* was identified as a prognostically relevant gene in a machine-learning-derived lysosome-dependent cell-death signature in neuroblastoma.^13^ In that study, *ST6GAL2* contributed to patient risk stratification and siRNA-mediated *ST6GAL2* knockdown reduced viability in five neuroblastoma cell lines. However, the relationship between ST6Gal2 expression, α2,6-sialylation, neuroblastoma clinical and transcriptional features, and other tumorigenic phenotypes remains unknown.

In this study, we investigate the clinical and functional significance of ST6Gal2 in neuroblastoma. We examine the association of *ST6GAL2* expression with patient outcome, clinicopathologic characteristics, and adrenergic and mesenchymal transcriptional signatures in independent publicly available neuroblastoma cohorts. We further generated stable *ST6GAL2* knockdown models to determine whether ST6Gal2 expression affects α2,6-sialylation and neuroblastoma cell proliferation, viability, migration, invasion, and wound closure. These studies define the relationship between ST6Gal2 and aggressive neuroblastoma phenotypes and provide preliminary evidence that ST6Gal2 knockdown reduces α2,6-sialylation in neuroblastoma.

## Methods

### Computational analyses of public neuroblastoma datasets

Association between *ST6GAL2* expression and patient survival was analyzed using the R2: Genomics Analysis and Visualization Platform (https://r2platform.com). We used the Sequencing Quality Control Consortium (SEQC) neuroblastoma cohort (n=498 primary tumors; GSE49710) with expression measured by Agilent 44K microarray (probe UKv4_A_32_P126157).^14^ Patients were stratified into high and low *ST6GAL2* expression groups using the median expression value as the cutoff (n=249 per group). The median was chosen *a priori* as a cutoff independent of outcome. Kaplan-Meier curves for overall survival and event-free survival were generated in R2 and differences between groups were assessed by the log-rank test. As sensitivity analyses, patients were also stratified by comparing the top and bottom quartiles of expression (n=125 per group) and by R2’s “scan” function, which tests all possible cutoffs with a minimum group size of 8 and applies a Bonferroni correction to account for multiple testing. Survival analyses were independently repeated in the Kocak neuroblastoma cohort (n=649 primary tumors; GSE45547).^15^ Patients with available survival data were dichotomized into high and low *ST6GAL2* expression groups using the median expression value (n=238 per group). Overall and event-free survival were estimated using the Kaplan-Meier method and differences between groups were assessed using the log-rank test.

Associations between *ST6GAL2* expression and clinicopathologic characteristics were evaluated in the SEQC cohort using R2. Expression was compared according to risk group, *MYCN* amplification status, stage, and sex. Patients were classified as high-risk or low/intermediate-risk according to the clinical risk annotation provided with the SEQC dataset, which was based on the German NB2004 risk stratification criteria.^16^ Stage was based on the International Neuroblastoma Staging System (INSS).^17^ Overall differences among multiple groups were assessed using one-way ANOVA. Pairwise comparisons were performed using Welch’s *t*-test with Bonferroni correction when necessary for multiple comparisons.

Gene expression data from the SEQC neuroblastoma cohort (n=498 primary tumors; GSE49710) were mapped from microarray probes to gene symbols using the GPL16876 platform annotation. Expression values for genes represented by multiple probes were averaged. Adrenergic (ADRN) and mesenchymal (MES) gene signatures were obtained from van Groningen *et al*.^5^ For each tumor, genes were ranked by expression and converted to percentile ranks. ADRN and MES signature scores were calculated as the mean percentile rank of signature genes represented on the array. Of the original signature genes described by van Groningen, 323/369 ADRN genes and 446/485 MES genes were represented. Associations between *ST6GAL2* expression and signature scores were assessed using Spearman rank correlation. The analysis was independently repeated in the Kocak neuroblastoma cohort (n=649 primary tumors; GSE45547) using the same gene-level averaging, percentile-rank signature scoring, and Spearman rank correlation procedure.

### Cell culture

Human neuroblastoma SK-N-AS and SK-N-BE(2) cells were maintained at 37°C in a humidified incubator containing 5% CO_2_. SK-N-AS cells were cultured in DMEM (Gibco, catalog no. 10-313-039) supplemented with 10% fetal bovine serum (FBS; Corning, catalog no. 35-011-CV), 1× non-essential amino acids (NEAA; HyClone, catalog no. SH3023801), 1× penicillin-streptomycin (HyClone, catalog no. SV30010), and 4 mM L-glutamine (HyClone, catalog no. SH3003401). SK-N-BE(2) cells were cultured in DMEM/F-12 (Gibco, catalog no. 21-331-020) supplemented with 10% FBS, 1× NEAA, 1× penicillin-streptomycin, and 2 mM L-glutamine. Cells were routinely monitored for mycoplasma and verified using STR fingerprinting.

### Generation of stable *ST6GAL2* knockdown cell lines

Stable *ST6GAL2* knockdown models were generated in SK-N-AS and SK-N-BE(2) neuroblastoma cells using purified lentiviral particles obtained from GeneCopoeia. For knockdown experiments, cells were transduced with lentiviral particles encoding three independent short hairpin RNAs (shRNA1-3) targeting human *ST6GAL2* (NM_001142351.2) in the psi-LVRU6GH vector containing a U6 promoter, eGFP, and a hygromycin-resistance cassette (GeneCopoeia, catalog no. LPP-HSH185245-LVRU6GH). Cells transduced with scrambled/non-targeting shRNA lentiviral particles served as negative controls (shControl; GeneCopoeia, catalog no. LP518). Cells were seeded in 6-well plates at 3×10^5^ cells per well and transduced when cells reached 70% confluency. Lentiviral particles were used at a multiplicity of infection (MOI) of 2 in the presence of polybrene (MilliporeSigma, catalog no. TR1003G) at a final concentration of 6 µg/mL. Viral stocks were 1×10^8^ TU/mL. Following transduction, cells were incubated at 37°C for 24 hours before replacement with fresh complete medium. Transduced populations were sorted by fluorescence-activated cell sorting (BD FACSAria III Cell Sorter, UAB Flow Cytometry and Single Cell Core Facility) into a pooled eGFP-positive subset and subsequently selected with hygromycin (Gibco, catalog no. 10-687-010) at 200 µg/mL for 10-14 days.

### Immunoblotting

Cells were harvested and lysed in RIPA buffer (Thermo Scientific, catalog no. J62524AE) supplemented with Halt protease/phosphatase inhibitor (Thermo Scientific, catalog no. 78440). Protein concentration was determined using Pierce BCA kit (Thermo Scientific, catalog no. 23250). A total of 20-50 µg of total protein per sample was separated by SDS-PAGE on premade gels (Bio-Rad Mini-PROTEAN TGX or GenScript SurePAGE) and transferred to PVDF membranes (Bio-Rad, catalog no. 1620177) using the Bio-Rad Trans-Blot Turbo transfer system. Membranes were blocked in 5% bovine serum albumin (BSA; Fisher BioReagents, catalog no. BP9706100) in Tris-Buffered Saline with Tween-20 (TBST; Bioland Scientific, catalog no. NC1530384) for 2 hours at room temperature and incubated with primary ST6Gal2 antibody (Proteintech, catalog no. 28367-1-AP) at 1:500 dilution in 5% BSA in TBST overnight at 4°C. Following washing, membranes were incubated with goat anti-rabbit secondary antibody (Invitrogen, catalog no. PI32460). Protein bands were detected using Luminata HRP substrate (Millipore, catalog nos. WBLUC0500, WBLUR0500, WBLUF0500) and imaged using the ChemiDoc imaging system (Bio-Rad). Primary antibody against β-actin (Invitrogen, catalog no. MA1140A488) was used at 1:5000 dilution in TBST for 2 hours at room temperature as a loading control. Protein bands were detected using fluorescence on the ChemiDoc imaging system (Bio-Rad).

### SNA flow cytometry

Cell surface α2,6-sialylation was assessed by flow cytometry using *Sambucus nigra* agglutinin (SNA). Cells were incubated with Cy5-conjugated SNA (Vector, catalog no. CL13051) at 10 µg/mL with 1% BSA in Hanks’ Balanced Salt Solution (HBSS; Thermo Scientific, catalog no. AAJ67771AP) for 1 hour on ice in the dark. Cells were then washed and resuspended in 1% BSA in HBSS for flow cytometric acquisition. Samples were acquired on a BD LSRFortessa (UAB Flow Cytometry and Single Cell Core Facility) and analyzed using FlowJo (Waters Biosciences). Cells were gated based on forward-and side-scatter, and SNA positivity was defined relative to unstained control. The percentage of SNA-positive cells was calculated for each condition.

### Cell proliferation and viability assays

Cell proliferation and viability were assessed in stable *ST6GAL2* knockdown neuroblastoma cell lines. Cells were seeded at 1.5×10^5^ cells per well and evaluated at 24, 48, 72, and 96 hours. At each time point, cell number and viability were determined by trypan blue exclusion using 0.4% trypan blue (Gibco, catalog no. 15250061) and cell counting was performed using a TC20 Automated Cell Counter (Bio-Rad). Equal volumes of cell suspension and trypan blue were mixed before counting. Viability was calculated as the percentage of viable cells among total counted cells.

### Transwell migration and invasion assays

Cell migration and invasion were assessed using Transwell inserts with 8 µm pores (Greiner Bio-One, catalog no. 662638). The underside of each insert was coated with 10 µg/mL type I collagen (Cultrex, catalog no. 3440-100-01) at 37°C. For invasion assays, the upper surface of the insert was additionally coated with 50 µL of Matrigel (Corning, catalog no. 354234) diluted to 1 mg/mL and allowed to polymerize for 4 hours at 37°C. For 24-well assays, 4×10^4^ cells were placed in 100 µL of serum-free medium in the upper chamber with 600 µL of 10% FBS-containing medium in the lower chamber to act as a chemoattractant. For migration assays, cells were allowed to migrate for 24 hours, whereas invasion assays were incubated for 48 hours. Non-migrated/non-invaded cells were removed from the upper surface of the membrane by gentle wiping. Cells on the underside of the membrane were fixed with 3% paraformaldehyde for 10 minutes. Inserts were stained with 0.5% crystal violet (Thermo Scientific, catalog no. 405830250) for 15 minutes, washed with PBS, dried, and imaged by light microscopy. The area of membrane covered by migrated or invaded cells was quantified in pixels using FIJI.^18^

### Scratch wound-healing assays

Cells for scratch wound-healing assays were seeded at 2.5×10^5^ cells per well in 12-well plates and grown to 80-90% confluence. A linear scratch was made in the monolayer using a sterile 200 µL pipette tip. Cells were washed with PBS to remove detached cells and replaced with standard growth medium. Images were acquired immediately after scratching and at subsequent time points. Wound area was quantified relative to the initial wound area using FIJI and percent wound closure was calculated.

### Statistical analysis

For analyses of public neuroblastoma datasets, Kaplan-Meier survival distributions were compared using log-rank tests. Overall differences in *ST6GAL2* expression among multiple clinicopathologic groups were assessed by one-way ANOVA with pairwise Welch tests and Bonferroni correction where indicated. The R2 cutoff-scan analysis incorporated Bonferroni correction for testing multiple candidate expression cutoffs. Associations between *ST6GAL2* expression and ADRN or MES signature scores were assessed using Spearman rank correlation.

Data from *in vitro* experiments are presented as mean ± SEM unless otherwise indicated. Each experiment except for SNA flow cytometry was performed in triplicate and technical replicates were averaged within each independent biological experiment before statistical analysis. For proliferation, viability, and scratch wound-healing assays, *ST6GAL2* knockdown conditions were compared with the shControl from the same biological experiment at individual time points using two-tailed *t*-tests. For Transwell migration and invasion assays, values from each independent biological experiment were normalized to shControl, which was assigned a value of 1.0, and normalized shRNA values were compared with 1.0 using two-tailed *t-*tests. A p value < 0.05 was considered statistically significant.

Experiments consisting of technical replicates from a single experiment, including the preliminary SNA flow cytometry, were analyzed descriptively and were not subjected to statistical testing. These data are presented as mean ± SD.

## Results

### *ST6GAL2* expression is associated with adverse clinical features and outcomes in neuroblastoma

*ST6GAL2* expression was associated with adverse clinical features in the SEQC neuroblastoma cohort. *ST6GAL2* expression was significantly higher in high-risk tumors (n=176) than in low/intermediate-risk tumors (n=322; p=9.53×10^−5^; **Figure 1A**). Consistent with this association, patients with high *ST6GAL2* expression had significantly worse overall and event-free survival than patients with low *ST6GAL2* expression (n=249 per group; p=4.34×10^−3^ and p=2.19×10^−3^, respectively; **Figure 1B-C**).

**Figure 1.**
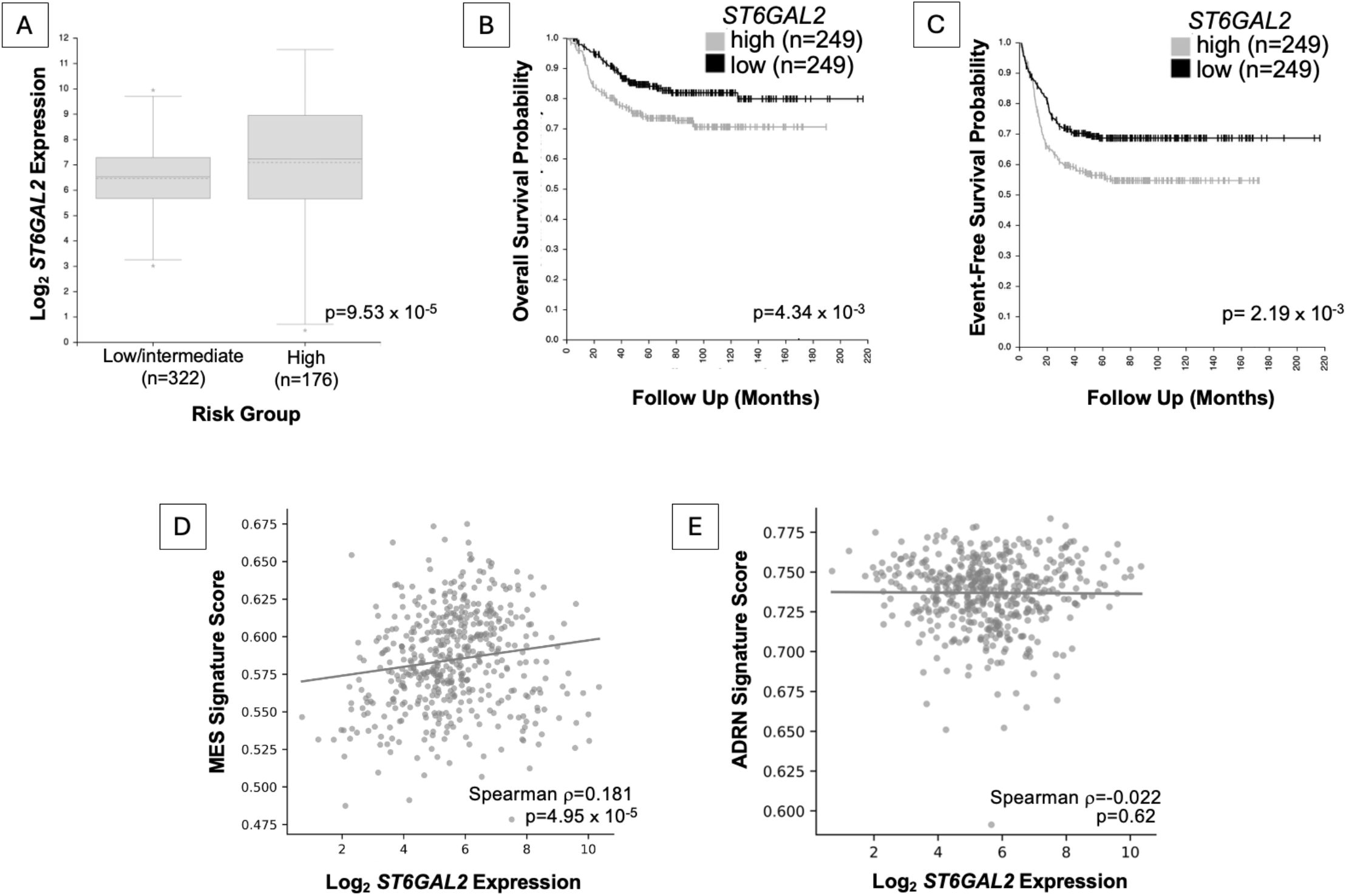
*ST6GAL2* expression is associated with clinically aggressive and mesenchymal-associated features in the SEQC neuroblastoma cohort. (A) *ST6GAL2* expression was higher in high-risk tumors (n=176) compared to low/intermediate-risk tumors (n=322; p=9.53×10^−5^). Kaplan-Meier analysis of (B) overall survival and (C) event-free survival in patients dichotomized into *ST6GAL2*-high (n=249) and *ST6GAL2*-low (n=249) groups by median expression value revealed a decrease in both overall survival and event-free survival in patients with high *ST6GAL2* expression (p=4.34×10^−3^ and p=2.19×10^−3^, respectively). Tick marks indicate censored observations. *ST6GAL2* expression was positively associated with the (D) van Groningen mesenchymal (MES) signature score (Spearman ρ=0.181, p=4.95×10^−5^) but was not significantly associated with the (E) van Groningen adrenergic (ADRN) signature score (Spearman ρ=−0.022, p=0.62). Each point represents one tumor. Straight lines indicate fitted trends for visualization, but associations were assessed using Spearman rank correlation.

To assess the robustness of the association between *ST6GAL2* expression and clinical outcome, we repeated the survival analyses using alternative expression cutoffs in the SEQC cohort. Patients in the highest quartile of *ST6GAL2* expression had significantly poorer overall survival (p=6.67×10^−3^) and event-free survival (p=6.13×10^−4^) than those in the lowest quartile (**Supplementary Figure 1A-B**). An outcome-based cutoff scan similarly identified significant associations between high *ST6GAL2* expression and poorer overall and event-free survival (p=5.11×10^−12^ and p=1.01×10^−7^, respectively; **Supplementary Figure 1C-D**). These sensitivity analyses support the association between elevated *ST6GAL2* expression and adverse outcome observed using the prespecified median cutoff.

We further examined associations between *ST6GAL2* expression and clinicopathologic features in the SEQC cohort. *ST6GAL2* expression was higher in *MYCN*-amplified than in *MYCN*-nonamplified tumors (p=1.44×10^−21^; **Supplementary Figure 2A**). Expression also differed significantly across INSS stages (p=3.04×10^−5^; **Supplementary Figure 2B**). Pairwise analyses demonstrated higher *ST6GAL2* expression in stage 4S tumors than in stages 1, 2, 3, and 4 after correction for multiple comparisons. *ST6GAL2* expression did not significantly differ by sex (p=0.71; **Supplementary Figure 2C**).

### *ST6GAL2* expression is associated with mesenchymal transcriptional features in the SEQC neuroblastoma cohort

To determine whether *ST6GAL2* expression was associated with established neuroblastoma cell states, we calculated van Groningen ADRN and MES signature scores in the SEQC cohort. *ST6GAL2* expression correlated positively with the MES signature score (Spearman ρ=0.181, p=4.95×10^−5^) but showed no significant association with the ADRN signature score (Spearman ρ=−0.022, p=0.62), suggesting that *ST6GAL2* expression is preferentially associated with mesenchymal-related transcriptional features (**Figure 1D-E**).

### Clinical and transcriptional associations of *ST6GAL2* are independently reproduced in the Kocak neuroblastoma cohort

We next assessed whether the clinical and transcriptional associations of *ST6GAL2* observed in the SEQC cohort were reproduced in the independent Kocak neuroblastoma cohort (GSE45547; n=649). Using the same median expression value cutoff as above, high *ST6GAL2* expression was again associated with poorer overall survival (n=238 per group; p=1.09×10^−4^) and event-free survival (n=238 per group; p=2.66×10^−4^; **Supplementary Figure 3A-B**). Consistent with the SEQC analysis, *ST6GAL2* expression was positively correlated with the MES signature score (Spearman ρ=0.204, p=1.51×10^−7^; **Supplementary Figure 3C**). A weaker positive correlation was also observed with the ADRN signature score (Spearman ρ=0.093, p=0.02; **Supplementary Figure 3D**). Together, these analyses independently reproduce the association of elevated *ST6GAL2* expression with adverse clinical outcome and mesenchymal-related transcriptional features.

### Stable knockdown of ST6Gal2 reduces α2,6-sialylation in neuroblastoma cells

Because *ST6GAL2* expression was associated with *MYCN* amplification in patient tumors, we used SK-N-AS (*MYCN*-nonamplified) and SK-N-BE(2) (*MYCN*-amplified) cells to evaluate ST6Gal2-associated phenotypes in both contexts. To establish neuroblastoma models with altered ST6Gal2 expression, we generated stable *ST6GAL2* knockdown (KD) cell lines using SK-N-AS and SK-N-BE(2) cells. Immunoblotting confirmed reduced ST6Gal2 protein expression following shRNA-mediated knockdown in both cell lines, with shRNA3 producing the greatest reduction in ST6Gal2 abundance in SK-N-AS cells and shRNA2 producing the greatest reduction in ST6Gal2 abundance in SK-N-BE(2) cells (**Figure 2A**). These shRNAs were therefore selected for subsequent phenotypic studies. To assess whether altered ST6Gal2 abundance was accompanied by changes in α2,6-sialylation, we measured cell surface binding of *Sambucus nigra* agglutinin (SNA), a lectin that preferentially binds terminal α2,6-linked sialic acid,^19^ by flow cytometry in SK-N-AS cells. SNA positivity was reduced with *ST6GAL2* knockdown, consistent with a role for ST6Gal2 in promoting α2,6-sialylation (**Figure 2B**).

**Figure 2.**
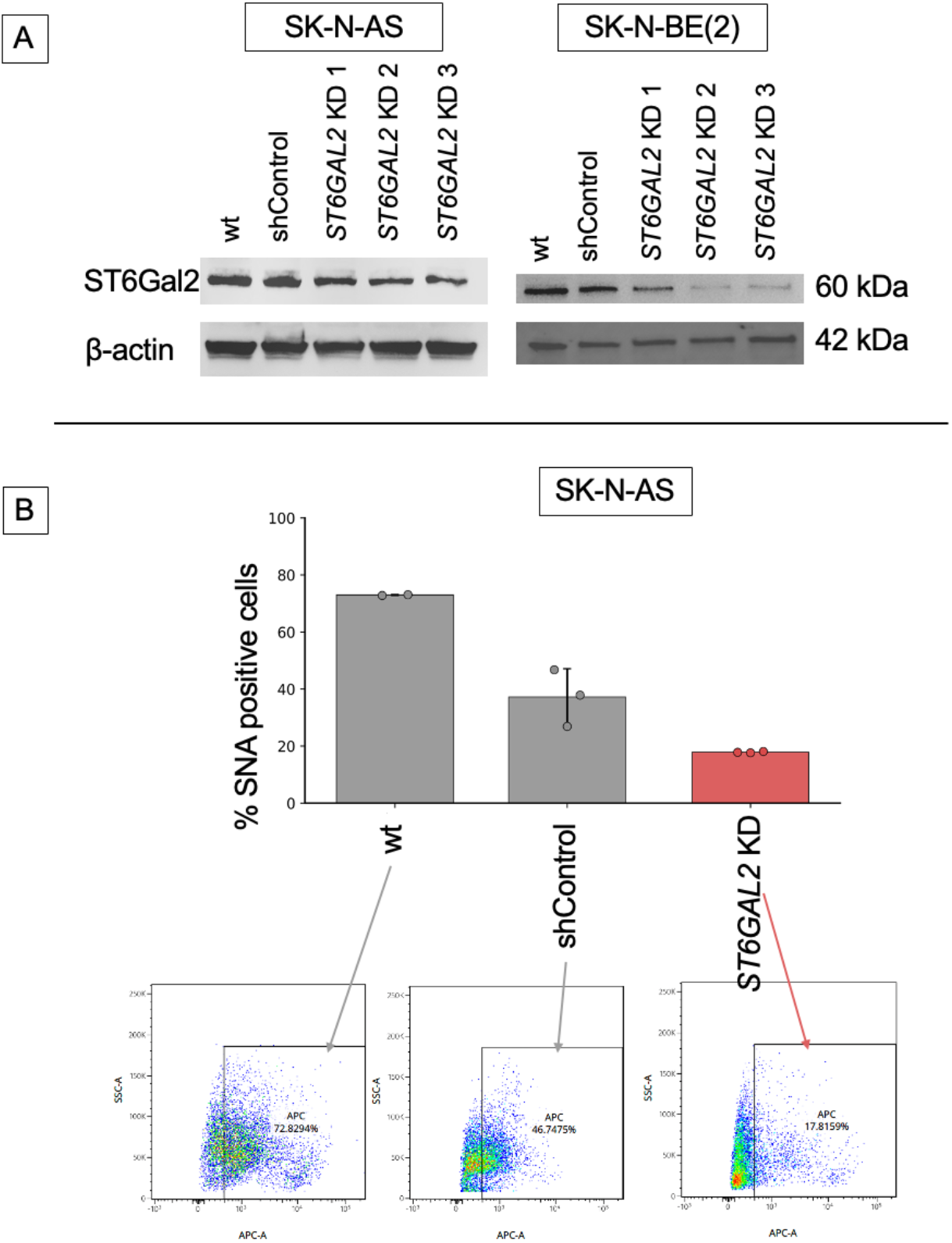
Stable *ST6GAL2* knockdown reduces α2,6-sialylation in neuroblastoma cells. (A) Representative immunoblots of ST6Gal2 expression in SK-N-AS and SK-N-BE(2) wild type (wt), non-targeting shRNA control (shControl), and stable *ST6GAL2* knockdown (KD) cells generated using three independent shRNAs (shRNA1-3). β-actin served as a loading control. shRNA3 produced the greatest reduction in ST6Gal2 protein expression in SK-N-AS cells whereas shRNA2 produced the greatest reduction in ST6Gal2 protein expression in SK-N-BE(2) cells. (B) SNA flow cytometric analysis of α2,6-sialylation in SK-N-AS cells. *ST6GAL2* knockdown showed a reduction in the proportion of SNA-positive cells. Representative flow cytometry plots are shown below the corresponding quantification. Bars represent mean ± SD of technical replicates from a single experiment.

### ST6Gal2 knockdown reduces neuroblastoma cell proliferation and viability

To determine whether ST6Gal2 contributes to neuroblastoma cell growth, proliferation was assessed over time in SK-N-AS and SK-N-BE(2) cells with *ST6GAL2* knockdown. In SK-N-AS cells, shControl cell counts increased from 0.15×10^6^ to 2.94 ± 0.08×10^6^ cells at 96 hours. *ST6GAL2* knockdown produced a reduction in cell proliferation, reaching only 1.29 ± 0.28×10^6^ cells at 96 hours (**Figure 3A**). There was significantly reduced cell proliferation with *ST6GAL2* knockdown at 48 hours (p=0.036) with trends toward reduced cell numbers at 72 hours (p=0.058) and 96 hours (p=0.077). In SK-N-BE(2) cells, shControl cells reached 1.35 ± 0.01×10^6^ cells at 96 hours compared to 0.69 ± 0.01×10^6^ cells with *ST6GAL2* knockdown (**Figure 3B**). *ST6GAL2* knockdown significantly reduced cell numbers relative to shControl at 48, 72, and 96 hours (p<0.01).

**Figure 3.**
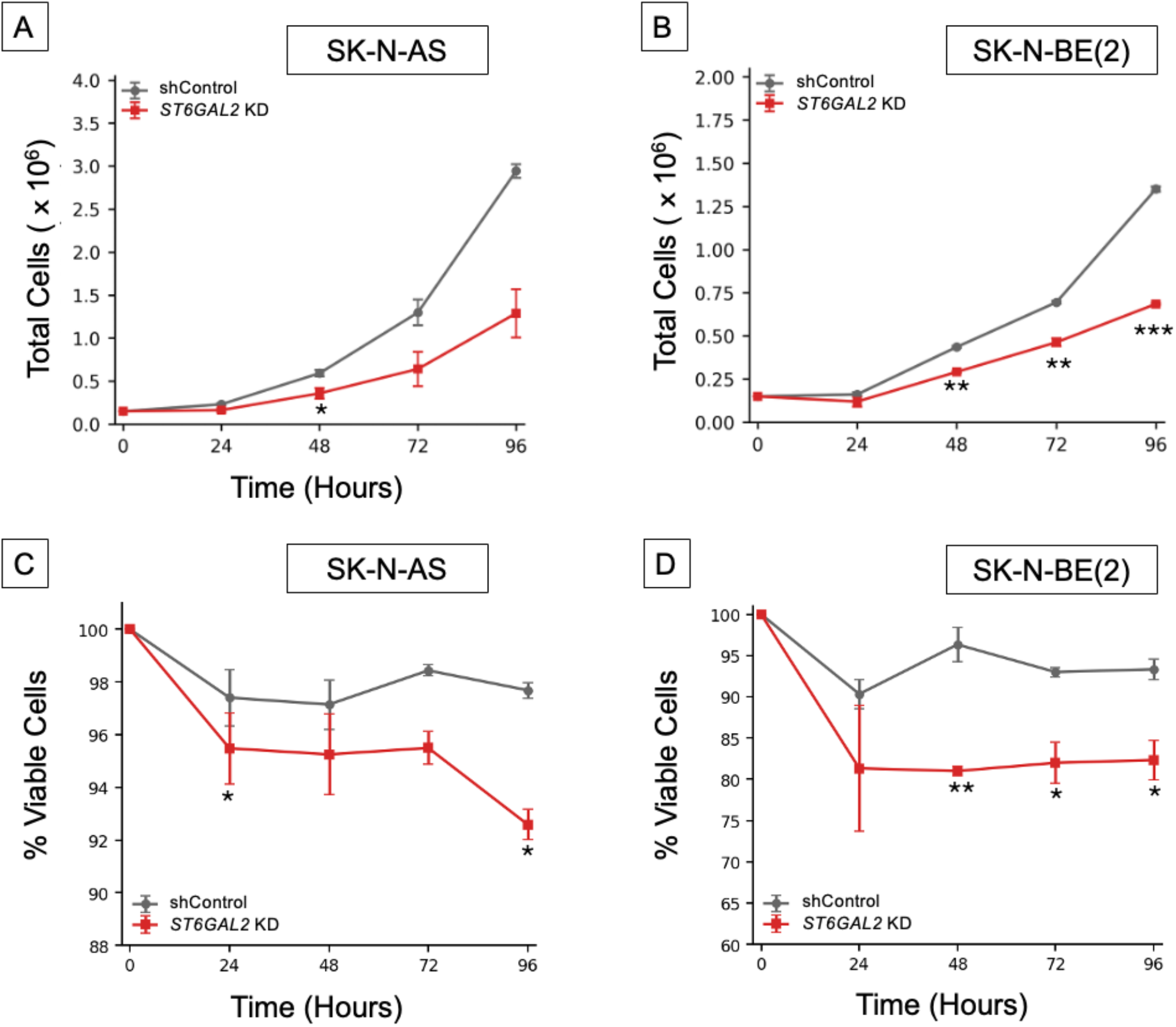
*ST6GAL2* knockdown reduces neuroblastoma proliferation and viability. Cell proliferation was assessed over time in (A) SK-N-AS and (B) SK-N-BE(2) cells stably expressing a non-targeting shRNA control (shControl) or *ST6GAL2*-targeting shRNA to achieve knockdown (*ST6GAL2* KD). Total viable cell numbers were determined at the indicated time points. Cell viability was assessed by trypan blue exclusion in (C) SK-N-AS and (D) SK-N-BE(2) cells. Viability at 0 hours was set to 100% for each condition. Experimental viability measurements were obtained beginning at 24 hours. Data are presented as mean ± SEM from independent biological experiments. Statistical comparisons between each knockdown condition and its matched shControl were performed at individual time points using two-tailed *t*-tests. *p<0.05, **p<0.01, ***p<0.001.

We next assessed cell viability by trypan blue exclusion (**Figure 3C-D**). In SK-N-AS cells, *ST6GAL2* knockdown produced a modest reduction in viability. At 96 hours, viability was 97.7 ± 0.3% in shControl cells versus 92.6 ± 0.6% in *ST6GAL2* knockdown cells. *ST6GAL2* knockdown demonstrated significantly reduced viability at 24 and 96 hours (p=0.019 and p=0.036, respectively), whereas differences at 48 and 72 hours did not reach statistical significance (p=0.093 and p=0.070, respectively). In SK-N-BE(2) cells, viability at 96 hours was 93.3 ± 1.2% in shControl cells compared with 82.3 ± 2.4% in *ST6GAL2* knockdown cells. *ST6GAL2* knockdown significantly reduced viability relative to shControl at 48, 72, and 96 hours (p<0.05). Together, these findings indicate that reduced ST6Gal2 expression impairs neuroblastoma cell growth and viability.

### ST6Gal2 knockdown impairs neuroblastoma cell migration, invasion, and wound closure

To determine whether ST6Gal2 contributes to neuroblastoma cell motility, Transwell migration and invasion assays were performed in SK-N-BE(2) cells with *ST6GAL2* knockdown. Relative to shControl cells, *ST6GAL2* knockdown reduced migration by 46% (p=0.040; **Figure 4A**). Similarly, *ST6GAL2* knockdown significantly reduced invasion by 66% relative to shControl (p=0.0047; **Figure 4B**).

**Figure 4.**
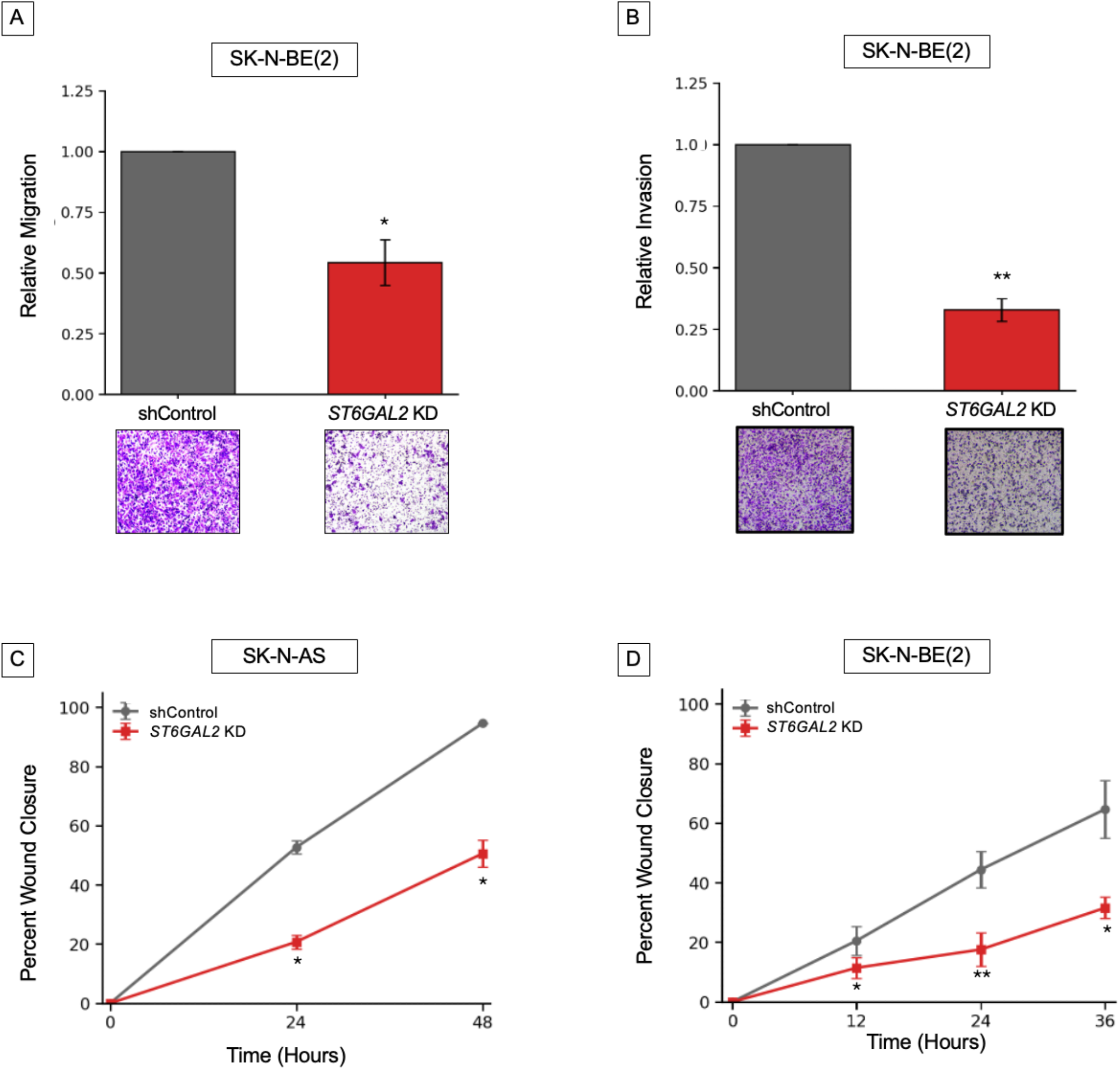
*ST6GAL2* knockdown reduces neuroblastoma migratory and invasive behavior. (A) Transwell migration in SK-N-BE(2) *ST6GAL2* knockdown (KD) cells was normalized to shControl within each independent biological experiment. (B) Transwell invasion in SK-N-BE(2) *ST6GAL2* knockdown (KD) cells was normalized to shControl within each independent biological experiment. Data are presented as mean ± SEM. Representative images of inserts are shown for each condition. Scratch wound-healing assays were performed in (C) SK-N-AS and (D) SK-N-BE(2) cells expressing shControl or shRNA targeting *ST6GAL2* to achieve knockdown (*ST6GAL2* KD). Percent wound closure was calculated relative to the initial wound area at 0 hours. Data are presented as mean ± SEM. *p<0.05, **p<0.01.

Scratch wound-healing assays were also used to assess cell migration in both SK-N-AS and SK-N-BE(2) cells (**Figure 4C-D**). In SK-N-AS cells, *ST6GAL2* knockdown showed reduced wound closure with 20.7% closure at 24 hours and 50.6% closure at 48 hours compared to 52.7% and 94.6% in shControl cells (p=0.020 and p=0.011, respectively). In SK-N-BE(2) cells, *ST6GAL2* knockdown significantly reduced wound closure compared to shControl at 12, 24, and 36 hours (p=0.043, p=0.0031, and p=0.034, respectively). Together, these findings support a role for ST6Gal2 in promoting neuroblastoma cell motility and invasive behavior.

## Discussion

In this study, we identify ST6Gal2 as a sialyltransferase associated with aggressive neuroblastoma features and provide functional evidence that ST6Gal2 contributes to α2,6-sialylation and tumorigenic behavior in neuroblastoma cells. In independent publicly available patient cohorts, elevated *ST6GAL2* expression was associated with inferior overall and event-free survival, high-risk disease, and *MYCN*-amplified tumors. Expression varied significantly across INSS stages. In parallel, *ST6GAL2* expression correlated positively with a mesenchymal transcriptional signature. Experimentally, ST6Gal2 knockdown decreased α2,6-sialylation and neuroblastoma cell proliferation, viability, migration, invasion, and wound closure. These findings extend the emerging association between ST6Gal2 and aggressive disease in neuroblastoma.

Our analysis independently demonstrates an association between high *ST6GAL2* expression and poor survival using prespecified median expression cutoffs, with additional sensitivity analyses using quartile-based and outcome-scanned cutoffs with the same findings. Importantly, our data also place *ST6GAL2* within established clinicopathologic features of aggressive neuroblastoma including high-risk classification and *MYCN* amplification. The relationship between *ST6GAL2* and INSS stage was not monotonic. Expression was highest in stage 4S tumors. Stage 4S is a distinct form of metastatic disease in infants, with dissemination confined to skin, liver, and bone marrow, that carries an excellent prognosis, frequently with spontaneous regression and minimal or no therapy.^17, 20^ The fact that expression is highest in this favorable group argues against simply interpreting *ST6GAL2* expression as a uniform marker of aggressiveness. Because ST6Gal2 is enriched in neural tissue, one possibility is that its expression in 4S disease reflects a developmental or differentiation-associated state of infant tumors rather than an aggressive phenotype, consistent with evidence that tumor-associated sialylation in neuroblastoma can track cell state and differentiation rather than aggressiveness alone.^21^ Alternatively, tumor sampling, cellular composition, or the small size of the 4S group may contribute.

The association between *ST6GAL2* expression and mesenchymal transcriptional features provides additional evidence of a link between ST6Gal2 and aggressive neuroblastoma biology. Neuroblastoma cells can be identified by adrenergic and mesenchymal transcriptional states and mesenchymal cells have been associated with greater invasiveness and therapeutic resistance.^5, 22, 23^ Mabe *et al*. demonstrated that adrenergic-to-mesenchymal transition alters expression of another sialyltransferase, ST8SIA1, resulting in reduced GD2 synthesis and resistance to anti-GD2 therapy.^24^ Therefore, neuroblastoma cell state is already known to be accompanied by remodeling of specific sialylation pathways. Our observation that *ST6GAL2* expression positively correlates with the mesenchymal signature raises the possibility that α2,6-sialylation represents another glycosylation event linked to neuroblastoma cell identity. However, the magnitude of the correlation in both cohorts reported here was modest, indicating that *ST6GAL2* expression should not be considered a surrogate marker for cell state. Instead, this indicates that elevated *ST6GAL2* expression tends to occur in tumors with mesenchymal-associated transcriptional features. Whether ST6Gal2 contributes directly to the establishment or maintenance of a particular neuroblastoma cell state cannot be determined from these data and will require dedicated mechanistic studies.

An additional finding of this study is that altering ST6Gal2 abundance was accompanied by changes in α2,6-sialylation. SNA staining decreased following *ST6GAL2* knockdown, providing preliminary evidence that ST6Gal2 contributes to α2,6-sialylation in neuroblastoma. This is consistent with the known enzymatic activity of ST6Gal2, which catalyzes α2,6-sialylation of terminal Galβ1,4GlcNAc-containing structures, although ST6Gal2 has more restricted substrate preferences than ST6Gal1.^8^ The best-characterized cancer-associated α2,6-sialyltransferase is ST6Gal1. Across multiple tumor types, ST6Gal1-mediated α2,6-sialylation has been linked to sustained proliferation, survival, plasticity, invasion, self-renewal and chemoresistance.^9-11, 25, 26^ These effects can arise through altered function of specific glycoprotein receptors and adhesion molecules. For example, α2,6-sialylation of β1 integrins can enhance adhesion, migration, and focal-adhesion signaling whereas altered EGFR α2,6-sialylation can influence receptor stability and promote chemoresistance.^27, 28^ Our findings suggest that the less widely expressed ST6Gal2 may contribute to analogous malignant phenotypes in neuroblastoma, but whether it acts through overlapping or distinct glycoprotein substrates remains unknown.

Emerging studies suggest that the effects of ST6Gal2 may be highly tissue dependent. In follicular thyroid carcinoma, ST6Gal2 was reported to promote proliferation, migration, invasion, angiogenesis, and tumor growth due to inactivation of Hippo signaling.^29^ However, reduced *ST6GAL2* expression has been associated with poorer prognosis in hepatocellular carcinoma,^30^ and a recent analysis of medulloblastoma data found an association between higher *ST6GAL2* expression and improved survival.^31^ Our findings place neuroblastoma among malignancies in which an increase in *ST6GAL2* expression is associated with aggressive behavior.

We found that ST6Gal2 knockdown impaired several phenotypes relevant to neuroblastoma progression including proliferation, viability, migration, and invasion. While cell viability was reduced, these effects were smaller and less uniform than the effects on proliferation. This indicates that the lower cell counts after ST6Gal2 knockdown may not be explained solely by loss of viability and likely also reflects altered proliferative capacity. However, the assays reported here do not distinguish effects on cell cycle progression, cell death, or other processes controlling cell accumulation.

Zi *et al*. recently reported reduced viability following transient *ST6GAL2* knockdown using siRNA in five neuroblastoma cell lines.^13^ Their findings exhibited a more pronounced effect on viability than what we observed with stable knockdown models, which may reflect differences in knockdown strategy, cell line context, assay methodology, or duration of gene suppression. The finding that ST6Gal2 knockdown reduced neuroblastoma cell motility and invasion is consistent with the existing literature linking altered sialylation to metastatic behaviors including changes in cell adhesion, integrin signaling, and receptor activation.^32^ In particular, ST6Gal1-mediated α2,6-sialylation has been associated with enhanced invasion and migration in several cancer types, in part through sialylation of integrins and growth factor receptors.^27^ The reduction in migration and invasion observed in this study is also directionally consistent with findings in follicular thyroid carcinoma, where *ST6GAL2* manipulation altered both migratory and invasive behavior.^29^

Several limitations must be considered. First, data from SNA flow cytometry assessing α2,6-sialylation are preliminary and derived from technical replicates of a single experiment, so these findings require confirmation in independent experiments and should be interpreted cautiously. Second, the clinical analyses are retrospective and based on bulk tumor transcriptomic datasets, so associations between *ST6GAL2* expression, mesenchymal signatures, and outcome cannot establish causality.

Despite these limitations, the study provides convergent clinical and experimental evidence supporting a role for ST6Gal2 in aggressive neuroblastoma biology. Elevated *ST6GAL2* expression was associated with adverse clinical outcome and mesenchymal-associated transcriptional features in patient cohorts while knockdown of ST6Gal2 reduced α2,6-sialylation and multiple tumorigenic phenotypes in neuroblastoma cells. These findings establish the biological relevance of ST6Gal2 in neuroblastoma.

## Acknowledgements

We thank the UAB Flow Cytometry and Single Cell Core Facility for providing instrumentation and services. The core facility is supported by the Center for AIDS Research (P30 AI027767) and the O’Neal Comprehensive Cancer Center (P30 CA013148), along with shared instrument grant S10OD032296.

## Author Contributions

Conceptualization: LS, KMH, LLS; Formal analysis: LS, LLS; Funding acquisition: KMH, LLS; Investigation: LS, PM, ML, SV, AR; Writing - Original Draft Preparation: LS, LLS; Writing - Review & Editing: LS, PM, ML, SV, AR, PJ, KMH, LLS.

## Funding

This work was supported by the American Pediatric Surgical Association Foundation (Jay Grosfeld, MD Scholar Grant, awarded to LLS) and the Kaul Pediatric Research Institute (KPRI) at Children’s of Alabama (awarded to LLS). The funders had no role in study design, data collection and analysis, decision to publish, or preparation of the manuscript.

## Notes

### Competing Interest Statement

The authors have declared no competing interest.

