## Supplementary Figures for "ST6Gal2 promotes α2,6-sialylation and aggressive phenotypes in neuroblastoma cells"

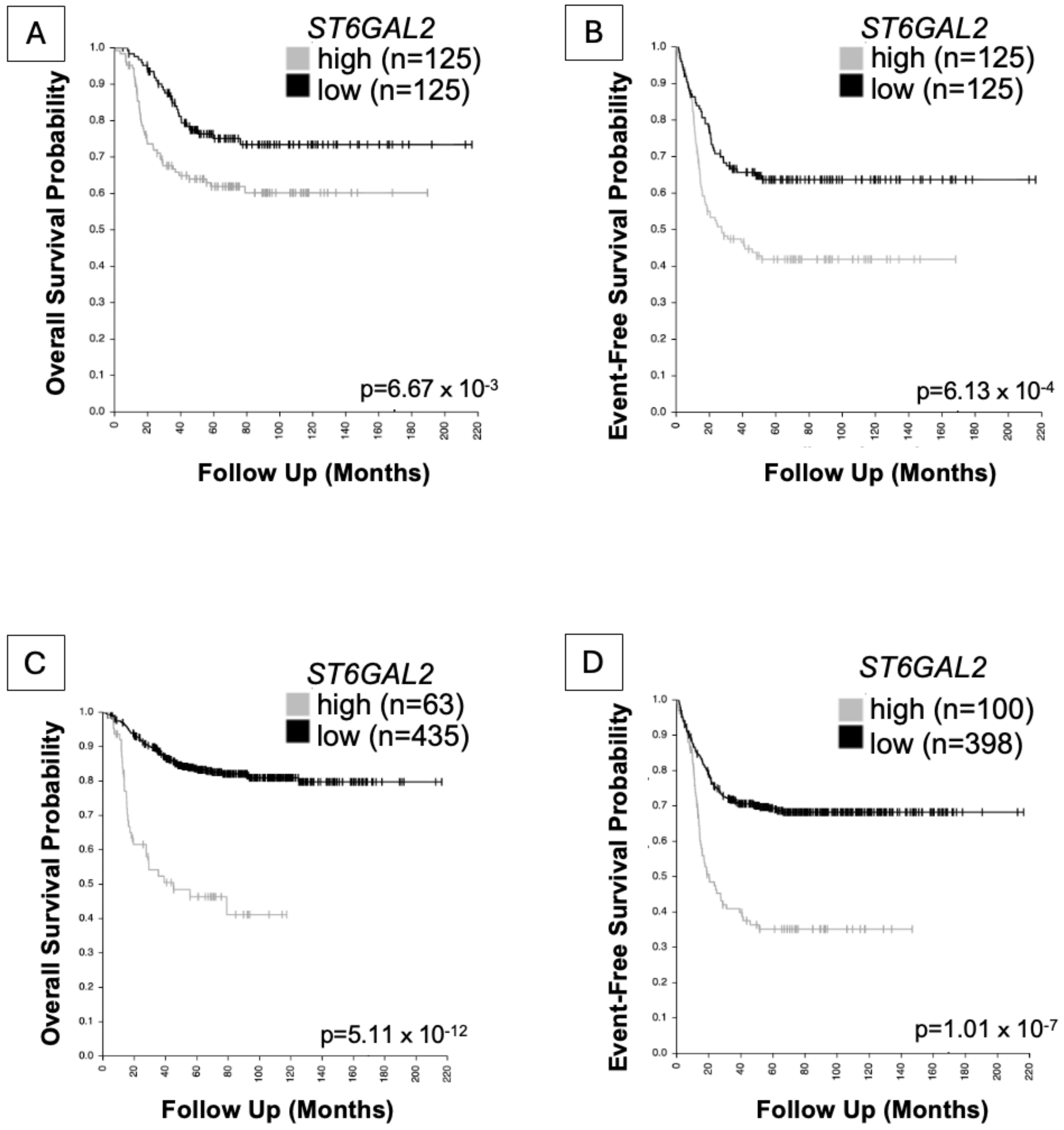


**Supplementary Figure 1.** Association between *ST6GAL2* expression and survival in the SEQC cohort remains significant with alternative expression cutoffs. Kaplan-Meier analysis of (A) overall and (B) event-free survival comparing patients in the highest and lowest quartiles of *ST6GAL2* expression (n=125 per group) revealed that high *ST6GAL2* expression was associated with poorer overall survival ($p=6.67\times10^{-3}$) and event-free survival ($p=6.13\times10^{-4}$). Kaplan-Meier analysis of (C) overall and (D) event-free survival using cutoffs identified by the R2 cutoff-scan procedure similarly showed that high *ST6GAL2* expression was associated with poorer overall survival (high n=63, low n=435; $p=5.11\times10^{-12}$) and event-free survival (high n=100, low n=398; $p=1.01\times10^{-7}$). Tick marks indicate censored observations.

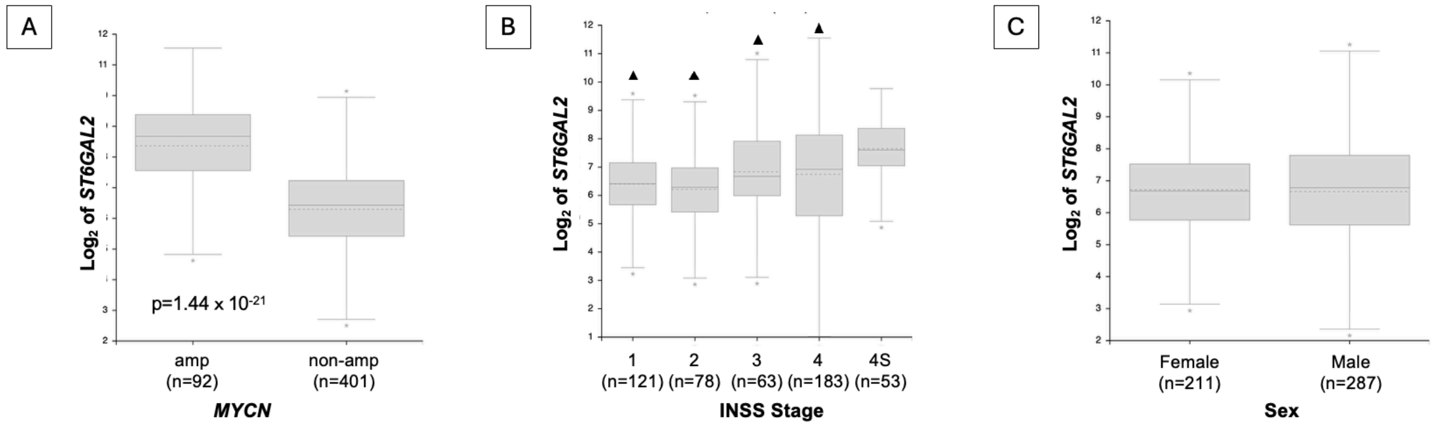


**Supplementary Figure 2.** Association of *ST6GAL2* expression with clinical features in the SEQC neuroblastoma cohort. (A) *ST6GAL2* expression was higher in *MYCN*-amplified neuroblastoma (n=92) than *MYCN*-nonamplified neuroblastoma (n=401; $p=1.44 \times 10^{-21}$). (B) *ST6GAL2* expression varied across INSS stages (stage 1 n=121, stage 2 n=78, stage 3 n=63, stage 4 n=183, stage 4S n=53; $p=3.04 \times 10^{-5}$). Triangles indicate stages with significantly different *ST6GAL2* expression compared with stage 4S in pairwise Welch tests after Bonferroni correction (adjusted $p<0.05$). (C) *ST6GAL2* expression did not differ by sex (female n=211, male n=287; p=0.71). Boxes indicate the interquartile range, with the median shown by the solid line and mean by the dashed line.

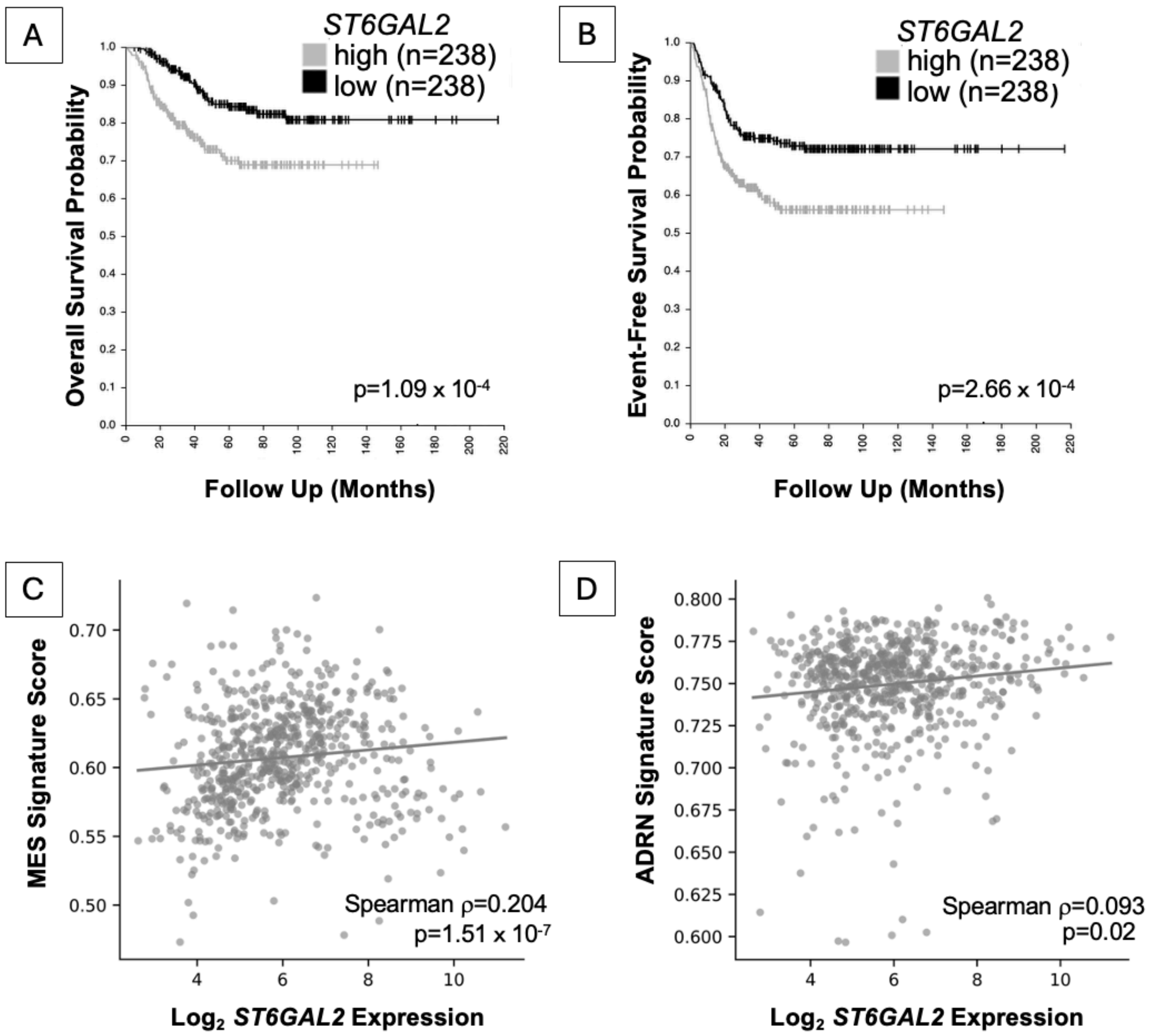


**Supplementary Figure 3.** Independent validation of clinical and transcriptional associations of *ST6GAL2* in the Kocak neuroblastoma cohort. Kaplan-Meier analysis of (A) overall survival and (B) event-free survival in patients dichotomized into *ST6GAL2*-high (n=238) and *ST6GAL2*-low (n=238) groups by median expression value revealed a decrease in both overall survival and event-free survival in patients with high *ST6GAL2* expression ($p=1.09 \times 10^{-4}$ and $p=2.66 \times 10^{-4}$, respectively). Tick marks indicate censored observations. *ST6GAL2* expression was positively associated with the (C) van Groningen mesenchymal (MES) signature score (Spearman $\rho=0.204$, $p=1.51 \times 10^{-7}$) and showed a weaker positive association with the (D) van Groningen adrenergic (ADRN) signature score (Spearman $\rho=0.093$, $p=0.02$). Each point represents one tumor. Straight lines indicate fitted trends for visualization, but associations were assessed using Spearman rank correlation.
